# Ultrastructural and Proteomic Signatures of Mechanoadaptive Fibroblast Remodeling across Microphysiological and Mesoscale Shear Platforms

**DOI:** 10.64898/2026.09.13.751238

**Authors:** Soyoun Min, Hyeon-Su Jin, Grahame Kidd, Emily Benson, Dong-Woo Lee, Hyun Jung Kim

## Abstract

Mechanobiological cues in the tissue microenvironment increasingly drive pathological fibroblast activation in inflammatory bowel disease (IBD), yet engineered platforms modeling this transition remain limited. Here, a microfluidic gut-on-a-chip microphysiological system and a mesofluidic rotary shaker are used to demonstrate that sustained fluid shear stress alone is necessary and sufficient to drive an irreversible, profibrotic phenotypic switch in primary normal human intestinal fibroblasts. Across both platforms, normal fibroblasts from small and large intestine reproducibly self-organize into three-dimensional (3D) multicellular aggregates within 72 h, independent of shear delivery format, indicating that the transition is governed by mechanical dose rather than device geometry. The resulting aggregates acquire robust α-smooth muscle actin (α-SMA) expression with aligned stress fibers, contrasting with the α-SMA-negative parental population. Scanning electron microscopy (SEM) resolves densely packed cellular microarchitecture embedded in a microfibrillar extracellular network, while ultrastructural serial block-face 3D EM reveals expansive intercellular spaces, stochastic fibrillar extrusions, and electron-dense cytoplasmic material at cell boundaries. Proteomic profiling confirms enrichment of core matrisome components, including collagen subtypes and matrix metalloproteinases. Together, these results establish fluid shear stress as a platform-independent, sufficient mechanical trigger for fibroblast-to-mechanoadaptive transition, positioning microphysiological shear platforms as tractable tools for modeling and targeting early fibrogenesis in IBD.

## 1. Introduction

Intestinal fibrosis represents a major unmet clinical challenge in inflammatory bowel disease (IBD), driving progressive tissue remodeling, luminal narrowing, and fibrostenotic complications that frequently require surgical intervention, particularly in Crohn’s disease (CD) [1, 2]. Despite significant advances in anti-inflammatory therapies, no approved anti-fibrotic treatments currently exist, reflecting a fundamental limitation in addressing fibrosis as a purely immune-mediated process. Increasing evidence indicates that fibrotic progression can persist independently of active inflammation, highlighting the importance of non-immune mechanisms, including aberrant mechanical signaling and tissue-level biomechanical remodeling, in sustaining disease progression [3]. Therefore, experimental systems capable of precisely controlling and interrogating mechanobiological cues are urgently needed to elucidate fibrosis initiation and accelerate the development of effective anti-fibrotic strategies.

In the intestinal mucosal microenvironment, epithelial barrier dysfunction represents an early pathological event that disrupts the spatial separation between luminal contents and the underlying stromal compartment. Loss of epithelial integrity exposes subepithelial fibroblasts to abnormal mechanical stimuli, including elevated fluid shear stress and altered extracellular matrix (ECM) tension, potentially triggering persistent fibroblast activation and maladaptive tissue remodeling [4]. Consistent with this concept, mechanically stimulated intestinal fibroblasts have been shown to undergo self-organization into ECM-rich three-dimensional (3D) multicellular aggregates exhibiting increased cellular stiffness comparable to that measured in human CD strictures [4]. These findings suggest that mechanical inputs alone can induce a disease-relevant fibroblast phenotype, providing a unique opportunity to engineer fibrosis-like tissue states independent of inflammatory stimulation. Importantly, establishing the reproducibility and scalability of this mechanically induced fibroblast transition across diverse cellular sources is a prerequisite for its development as a translational tissue engineered platform. Furthermore, resolving the 3D cellular organization and macromolecular signatures of these mechanically induced aggregates may enable identification of early fibrotic states and establish a tractable system for mechanistic investigation and therapeutic evaluation.

However, several critical limitations currently restrict the broader application of mechanically induced fibrosis models. Previous studies have primarily relied on a single mechanical stimulation format, leaving unresolved whether fibroblast aggregation represents a generalizable mechanoadaptive response or a phenomenon specific to a particular experimental configuration. Moreover, the ultrastructural organization, spatial architecture, and macromolecular composition underlying this mechanically transformed fibroblast state remain poorly characterized. Addressing these limitations requires an integrated engineering framework that combines controllable mechanical stimulation with high-resolution structural and molecular analysis to define how physical cues reshape fibroblast organization and function. Such a platform would not only provide mechanistic insight into fibrosis initiation but also establish a scalable foundation for patient-specific disease modeling and therapeutic screening.

Here, we establish a cross-platform engineering framework to determine whether fluid shear stress represents a transferable design parameter for inducing fibrosis-relevant fibroblast remodeling. Mechanically induced 3D fibroblast aggregates were generated using two mechanically distinct platforms: a microfluidic Gut-on-a-Chip microphysiological system (MPS) [4–8] and a mesoscale rotary shaker platform, enabling assessment of the reproducibility, scalability, and device independence of shear-driven tissue organization. We subsequently characterized these mechanically transformed aggregates using complementary structural and molecular approaches, including conventional scanning electron microscopy (SEM) and serial block-face SEM (SBF-SEM) to resolve 3D cellular and fibrillar architectures, together with proteomic profiling to identify ECM and cytoskeletal signatures associated with mechanical adaptation. Collectively, this study establishes fluid shear stress as a controllable bioengineering input for generating fibrosis-relevant multicellular architectures and provides a scalable platform for dissecting mechanobiological mechanisms underlying intestinal fibrosis. This work further positions mechanically engineered fibroblast aggregates as a translational model system for investigating fibrotic remodeling and evaluating next-generation anti-fibrotic interventions.

## 2. Results

### 2.1 Fluid shear stress induces irreversible 3D aggregation of normal intestinal fibroblasts

Consistent with our previous study [4], sustained fluid shear stress (0.0014 dyne cm^-2^) induced progressive remodeling of a shear-vulnerable normal intestinal fibroblast (nFib) monolayer cultured in the upper channel of a Gut-on-a-chip, resulting in the formation of hemispherical 3D multicellular aggregates on a polydimethylsiloxane (PDMS) membrane after approximately 160 h (Figure 1A). To evaluate whether this shear-induced aggregation was culture platform-independent, we reproduced the mechanical stimulation using a conventional rotary shaker in which nFib monolayers prepared in a 6-well plate were cultured on the identical PDMS membrane substrate (Figure 1B). Despite substantial differences in fluid dynamics and absolute shear magnitude (approximately 0.5-1 dyne cm^-2^), rotary-induced shear similarly disrupted the monolayer, formed cell clumps by 32 h, and rapidly generated robust 3D fibroblast aggregates within 48 h. These findings demonstrate that shear-driven aggregation is reproducible across mechanically distinct culture systems and is not dependent on a specific device configuration. Confocal immunofluorescence imaging further revealed a pronounced transition from a flattened 2D monolayer to compact 3D aggregates characterized by significantly increased cellular height (2.3-fold, *p*<0.0001) and marked upregulation of the profibrotic markers α-SMA and collagen I (COL I) (20.0- and 2.0-fold, respectively; *p*<0.002; Figures 1C and S1). Consistent with their profibrotic phenotype, 3D aggregates exhibited significantly higher specific pan-matrix metalloproteinases (MMP) enzymatic activity than the 2D monolayer fibroblasts (1.3-fold, *p*<0.01), suggesting enhanced ECM remodeling. Collectively, these results establish sustained fluid shear stress as a robust mechanical cue that drives fibroblast self-organization into contractile, profibrotic multicellular aggregates across both microfluidic and mesofluidic culture platforms.

**Figure 1.**
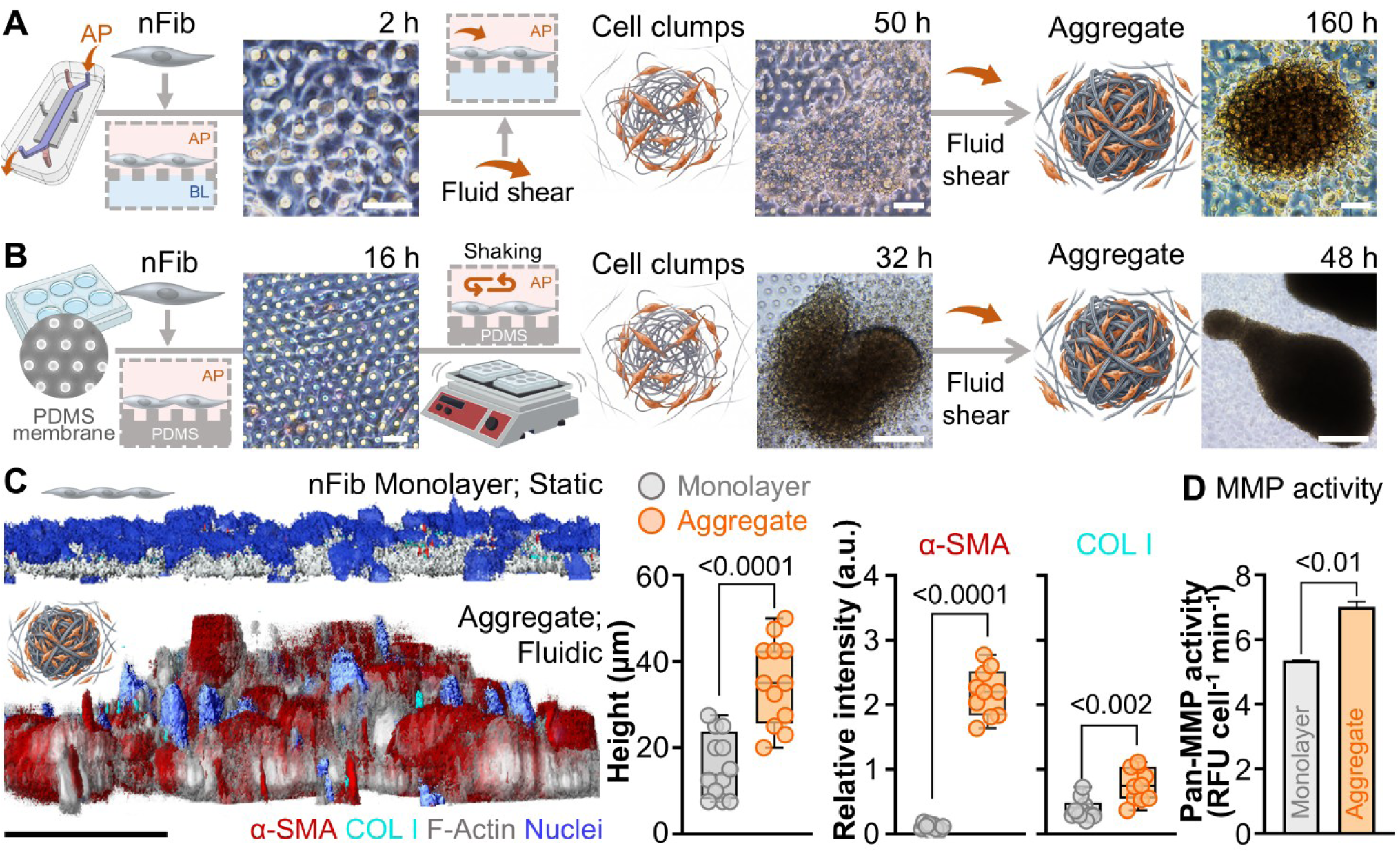
Fluid shear drives fibroblast-to-myofibroblast transition and ECM deposition across microfluidic and mesofluidic culture platforms. (A) Schematic of the microfluidic Gut-on-a-chip platform. Normal colonic fibroblasts (nFib) seeded on the apical (AP) channel surface adhere as a monolayer (2 h), self-assemble into cell clumps under fluid shear (50 h), and consolidate into a compact 3D aggregate by 160 h. BL, basolateral channel. (B) Schematic of the mesofluidic platform. nFib cells seeded on a PDMS membrane in a 6-well format (16 h) form cell clumps under orbital shaking-induced shear (32 h) and mature into aggregates by 48 h. Representative phase-contrast images are shown above each corresponding timepoint. Bars, 50 µm. (C) Volumetric confocal reconstruction of static nFib monolayers vs. fluidically stimulated aggregates immunostained for α-SMA (red), COL I (cyan), F-actin (grey), and nuclei (blue), illustrating the marked increase in tissue height and myofibroblast marker expression under shear. Quantification confirms significantly greater aggregate height, α-SMA intensity, and COL I intensity relative to static monolayer controls. (D) Pan-MMP activity is significantly elevated in fluidically stimulated aggregates compared with static monolayers, consistent with enhanced ECM turnover accompanying fibroblast-to-myofibroblast transition. Data are presented as mean ± SEM. Statistical significance was determined using an unpaired two-tailed Student’s *t*-test (C) or Welch’s *t*-test (D).

### 2.2 Ultrastructural analysis reveals a stacked cellular microarchitecture with a dense fibrillar network

Scanning electron microscopy revealed that fibroblast-derived 3D aggregates formed compact, irregularly lobulated structures measuring approximately 80-150 µm in diameter (Figure 2, Aggregate). Despite their compact morphology, the aggregates displayed substantial ultrastructural heterogeneity. High-magnification images showed that the aggregate surface was enmeshed in a dense, interwoven fibrillar network that bridged neighboring cellular domains (Figure 2A-D).

**Figure 2.**
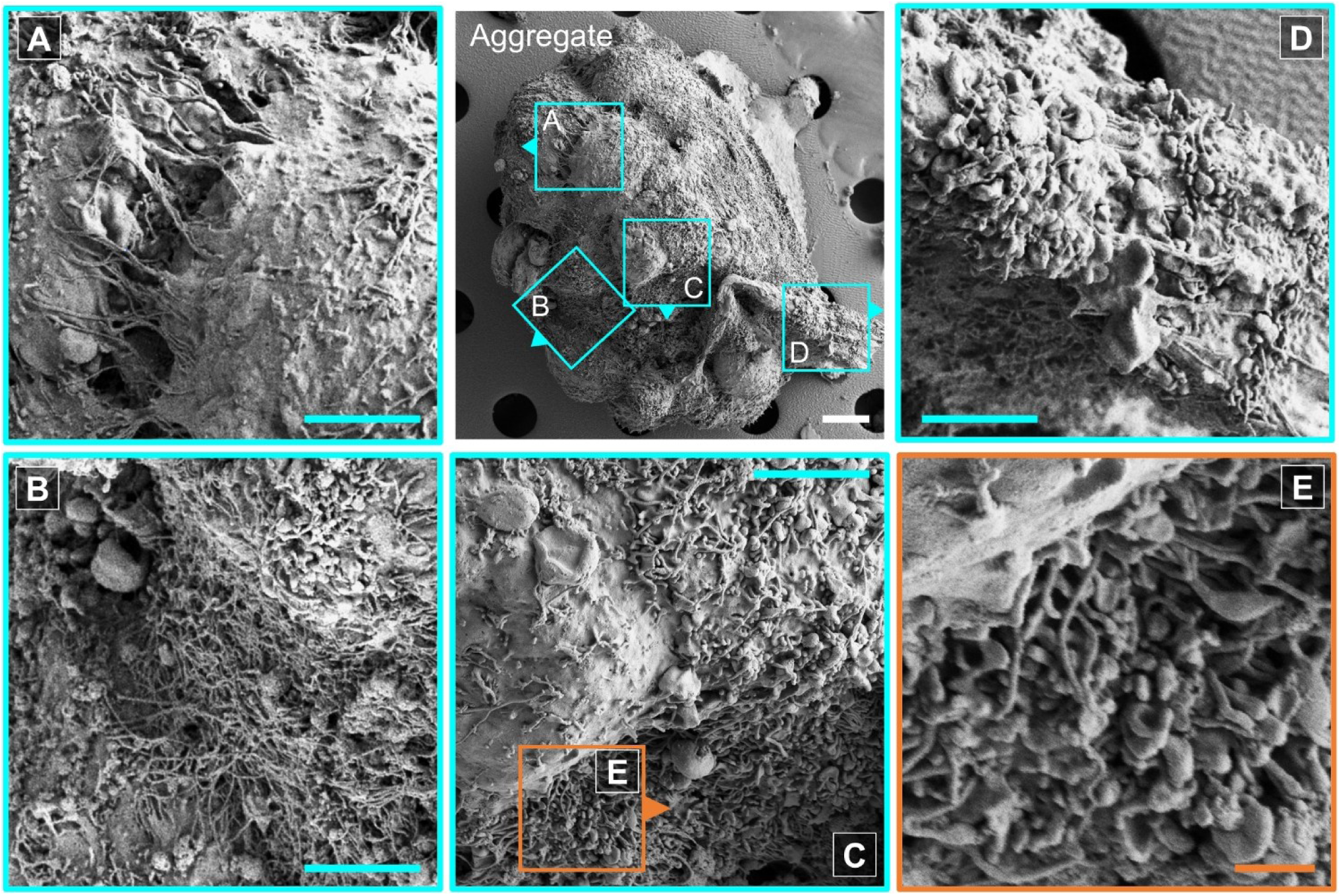
Scanning electron microscopy reveals a heterogeneous, fibril-rich surface ultrastructure across the shear-induced fibroblast aggregate. Center top: low-magnification SEM overview of an intact aggregate showing an irregular, lobulated surface topography. Boxed regions (2A-D, cyan) indicate areas selected for higher-magnification imaging. (A) Region corresponding to a peripheral lobule, showing a dense meshwork of extracellular fibrils interspersed with globular surface deposits and underlying cellular contours. (B) Region showing a transitional zone between a fibril-sparse, membrane-smooth surface (upper left) and a dense fibrillar network (lower right), illustrating spatial heterogeneity in matrix deposition across the aggregate surface. (C) Region encompassing a cell-dense protruding surface domain with an underlying fibrillar substrate; the boxed area (E, orange) marks a subregion selected for further magnification. (D) Region showing a cluster of closely apposed cell-surface protrusions with fine filamentous processes extending into the surrounding fibrillar matrix. (E) Higher-magnification view of the boxed region in (C), resolving individual interdigitating cell-surface processes embedded within a dense, porous fibrillar network. Bars in white, 10 µm; in cyan, 5 µm; in orange, 1 µm.

Quantitative morphometric analysis demonstrated distinct fibril populations within the same aggregate. Filaments in Figures 2A and 2B were relatively thin and uniform, with mean apparent widths of 0.130±0.014 µm (median, 0.127 µm; *n*=20) and 0.119±0.011 µm (median, 0.110 µm; *n*=20), respectively. In contrast, fibrils in Figures 2C and 2D appeared bundled or braided and exhibited significantly greater apparent widths (0.207±0.028 µm and 0.246±0.029 µm, respectively) together with broader size distributions (coefficient of variation, CV: 59.8% and 52.1%, respectively), distinguishing these bundled structures from the more uniform fibrillar population. At the highest magnification (Figure 2E), individual filaments spanned several micrometers and connected rounded nodular protrusions on the aggregate surface. These protrusions, morphologically resembling microvillus-like blebs, measured 0.323±0.026 µm in diameter (median, 0.295 µm; *n*=20). The extensive filamentous interconnections suggest that the aggregate surface is mechanically integrated through a continuous fibrillar scaffold.

This ultrastructural transition was consistently reproduced in fibroblasts isolated from multiple sources. Tissue-derived normal colonic fibroblasts (Figures 3A, B) and commercially obtained primary human colonic fibroblasts (Figures 3D, E), cultured on the same PDMS membrane under mesofluidic stimulation for 48 h, both formed compact 3D aggregates with diameters at 149.00±10.25 µm and 135.87±9.09 µm, respectively (Figure S2). Similar to small intestinal fibroblasts, both colonic fibroblast populations exhibited significantly higher specific pan-MMP activity than their corresponding 2D monolayers (1.43-fold and 1.39-fold increases, respectively), indicating enhanced MMP-dependent ECM remodeling during aggregate formation. SEM further confirmed that the dense fibrillar surface architecture was conserved across fibroblasts from both the small and large intestine, demonstrating that this mechanoadaptive transition is a robust and reproducible response to chronic fluidic stimulation, independent of anatomical origin.

**Figure 3.**
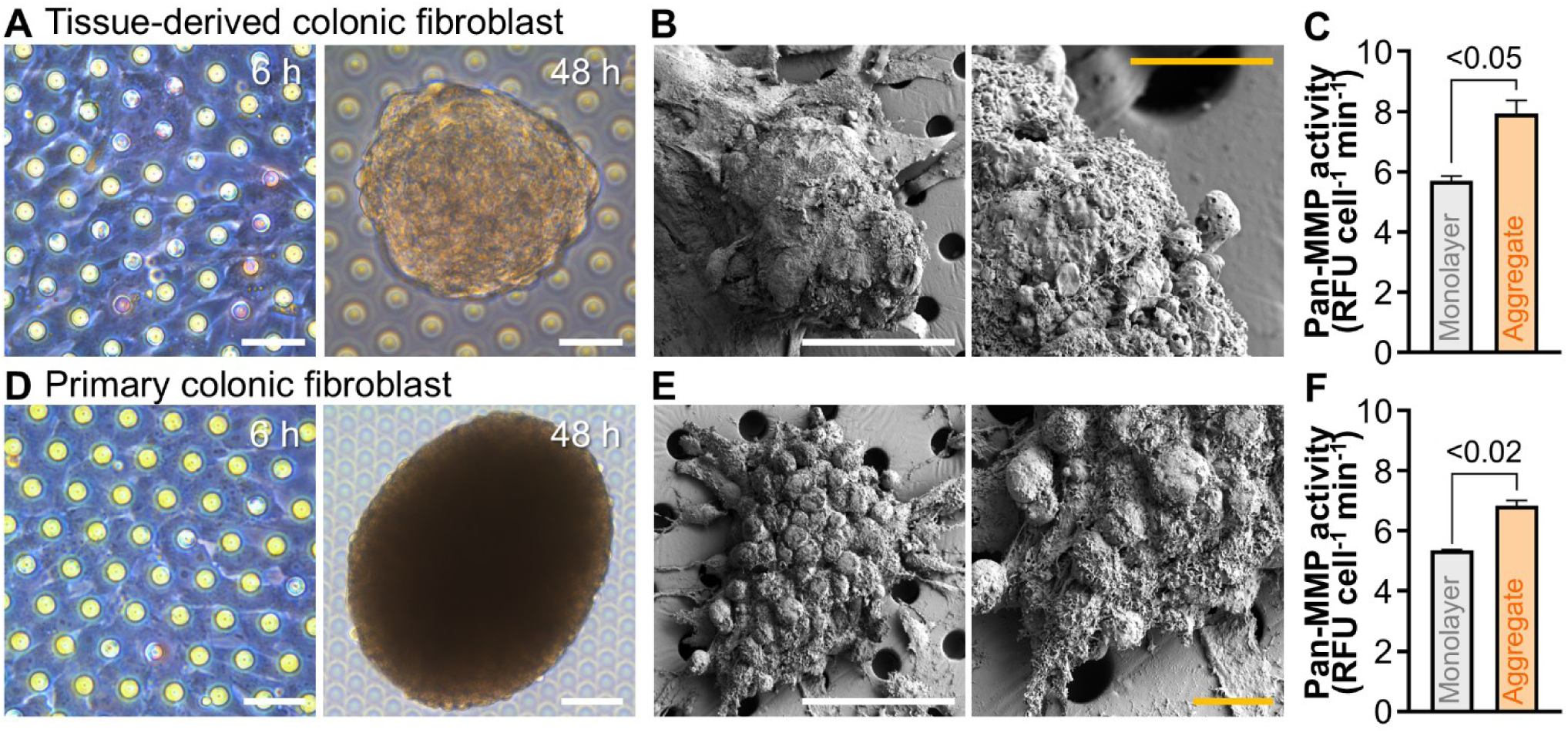
Cross-validation of shear-induced aggregate formation across two independent colonic fibroblast sources. (A) Tissue-derived colonic fibroblasts seeded on the mesofluidic PDMS membrane appear as an adherent monolayer at 6 h and consolidate into a compact 3D aggregate by 48 h (phase-contrast). (B) Representative SEM images of a tissue-derived fibroblast aggregate at two magnifications, showing a lobulated, fibril-rich surface topography consistent with the ultrastructural features described in Figure 2. (C) Pan-MMP activity is significantly elevated in tissue-derived fibroblast aggregates relative to static monolayer controls. (D) Primary colonic fibroblasts seeded under identical mesofluidic conditions similarly form an adherent monolayer at 6 h and mature into a compact 3D aggregate by 48 h. (E) Representative SEM images of a primary fibroblast aggregate at two magnifications, reproducing the dense, textured surface morphology observed in the tissue-derived model. (F) Pan-MMP activity is significantly elevated in primary fibroblast aggregates relative to 2D monolayer controls (*p*<0.02). Bars in white, 50 µm; in orange, 10 µm. Statistical significance was determined using Welch’s *t*-test.

### 2.3 Serial block-face SEM uncovers stochastic intercellular voids and heterogeneous fibrillar extrusion in aggregates

Serial block-face scanning electron microscopy (SBF-SEM) was used to resolve the 3D ultrastructure of fibroblast aggregates through automated serial nanometer-scale sectioning and image reconstruction (Figure 4A). The reconstructed aggregates comprised morphologically heterogeneous fibroblasts, ranging from elongated spindle-shaped to stellate style cells (Figures 4A middle and S3). Approximately half of the cells appeared ultrastructurally intact, containing a single oblate-to-rounded nucleus and intact organelles, whereas the remainder exhibited varying degrees of degeneration, including plasma membrane disruption or partial cellular fragmentation. Intact cells contained elongated mitochondria measuring approximately 0.2-0.5 µm in diameter and 1-10 µm in length, with occasional branching. The mitochondria displayed a homogeneous matrix, well-defined cristae, and minimal intracristal swelling, consistent with preserved structural integrity. 3D reconstruction further revealed an interconnected network of extracellular void spaces distributed throughout the aggregate and enriched near the periphery (Figure 4Aa, Aa’, demarcated violet region). These intercellular spaces provide structural accommodation for the extracellular fibrillar matrix while facilitating nutrient diffusion and metabolic waste exchange in the aggregate.

**Figure 4.**
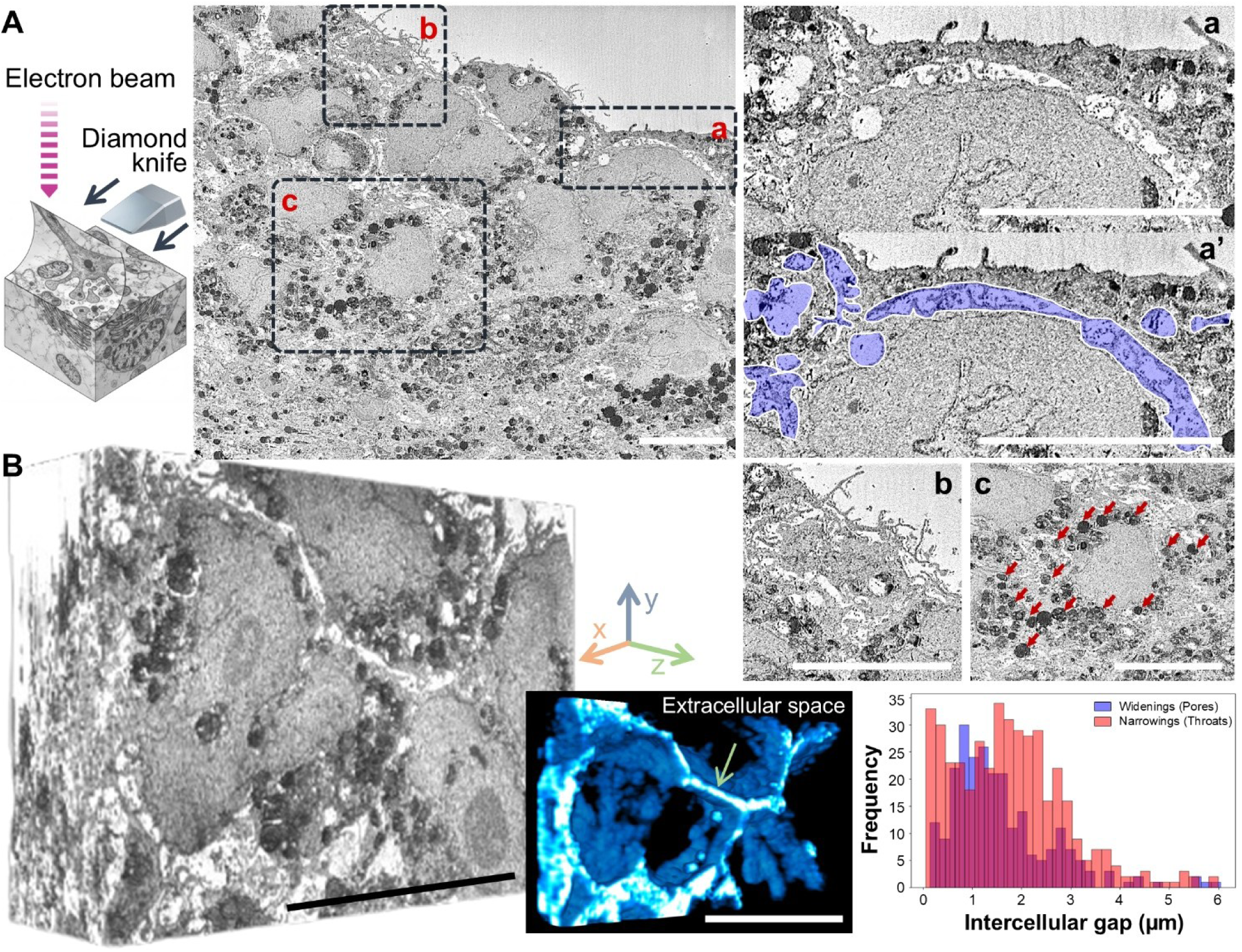
Serial block-face SEM reveals a 3D interconnected extracellular channel network with distinct pore–throat architecture within fibroblast aggregates. (A) Workflow of SBF-SEM, in which sequential ultrathin sectioning of a resin-embedded block face by an in-chamber diamond knife enables automated acquisition of volumetric electron microscopy image stacks. Representative block-face image showing interdigitating fibroblasts within the aggregate; boxed regions (a-c, dashed) are enlarged. (Aa) High-magnification view of a narrow extracellular channel enclosed by adjacent cell membranes. (Aa’) Segmentation of the extracellular space (purple overlay) used for morphometric analysis. (Ab) Interdigitating, organelle-poor cellular processes extending into the extracellular compartment. (Ac) Membrane-bound electron-dense bodies (red arrows) in the perinuclear/basal cytoplasm, consistent with lysosomal or autophagolysosomal structures. (B) 3D reconstruction of the aligned SBF-SEM stack demonstrating a continuous, tortuous extracellular channel network extending throughout the fibroblast aggregate (left). Segmented rendering of the extracellular space (cyan) highlights localized channel expansions (arrow) (middle). Histogram shows the distribution of extracellular gap widths, distinguishing pore-like widenings (blue) and narrow throat regions (red), revealing a predominantly sub-2 µm throat architecture with discrete larger pore domains (right). Bars, 10 µm.

Consistent with surface SEM observations (Figure 2), SBF-SEM confirmed an extensive meshwork extending both between neighboring cells and beyond the aggregate surface (Figure 4Ab). Numerous slender, organelle-poor cellular processes (0.3-0.5 µm in diameter and 5-20 µm in length) projected from the cell surface and frequently interdigitated with processes from adjacent cells. These projections generated a process-rich intercellular compartment approximately 1-5 µm wide, through which extracellular fibrils traversed the aggregate. Nearly all cells contained spherical membrane-bound electron-dense bodies (0.723±0.052; CV 39.1%) that were predominantly localized in the basal or perinuclear cytoplasm and, in some cells, immediately beneath the plasma membrane (Figure 4Ac, arrows). Phagosome-like vesicular structures containing heterogeneous membranous material were also frequently observed, indicating active intracellular membrane remodeling. Segmentation and 3D rendering of the reconstructed volume (Figure 4B, left panel; Movie S1) demonstrated that the intercellular compartment forms a continuous, branching channel network rather than isolated voids (Figure 4B, middle panel; Movie S2). Quantitative morphometric analysis further showed that the network is dominated by narrow “throat” segments interconnected by less frequent, locally expanded “pore” regions (Figure 4B, right), revealing a highly heterogeneous yet spatially continuous extracellular architecture within the fibroblast aggregate.

### 2.4 Proteomic profiling identifies enrichment of core ECM and cytoskeletal proteins in mechanically induced aggregates

Finally, we performed label-free quantitative proteomics on the conditioned media of the normal human intestinal fibroblasts (nFib) and the mechanoadaptive 3D aggregates (Agg) to define the extracellular protein landscape of mechanoadaptive aggregates. The nFib cells served as a reference representing the baseline secretory profile of non-activated fibroblasts under standard 2D culture conditions. Following common-protein median normalization (Figure S3A) and reproducibility filtering (≥2/3 replicates; Figure S3B), a total of 1,590 and 2,229 proteins were reproducibly detected in the Agg and nFib secretomes, respectively, with 1,390 proteins shared between the two conditions (Figure 5A). We annotated all detected proteins against the human matrisome using MatrisomeDB 2.0 [9] to systematically identify ECM-directed secretory activity. Matrisome proteins accounted for a substantial proportion of the total Agg secretome by relative abundance (38.22%), confirming the ECM-enriched character of the Agg secretory profile. Comparison of the Agg-specific and nFib-specific protein pools revealed differences in matrisome category distribution (Figure 5A and B). Among the 200 Agg-specific proteins, 41 (20.5%) were matrisome-annotated, compared with 44 of 839 (5.2%) nFib-specific proteins, indicating that, among proteins detected exclusively in each condition, those unique to Agg were disproportionately ECM-related (Figure 5A). Category-level comparison showed Agg-exclusive matrisome proteins skewed toward Secreted Factors (*n*=16; INS, PDGFD, S100A4), ECM Regulators (*n*=14; SERPINB9, SERPINF2, F2), ECM Glycoproteins (*n*=10; FBN3, FGB, FGA), Proteoglycan (*n*=2; SRGN, SPOCK1), and Collagens (*n*=2; COL8A1, COL15A1), whereas nFib-exclusive matrisome proteins were enriched for ECM Regulators (*n*=19; A2ML1, ADAMDEC1, CELA2A) and ECM-affiliated Proteins (*n*=7; ANXA3, SEMA6D, LMAN1) (Figure 5B).

**Figure 5.**
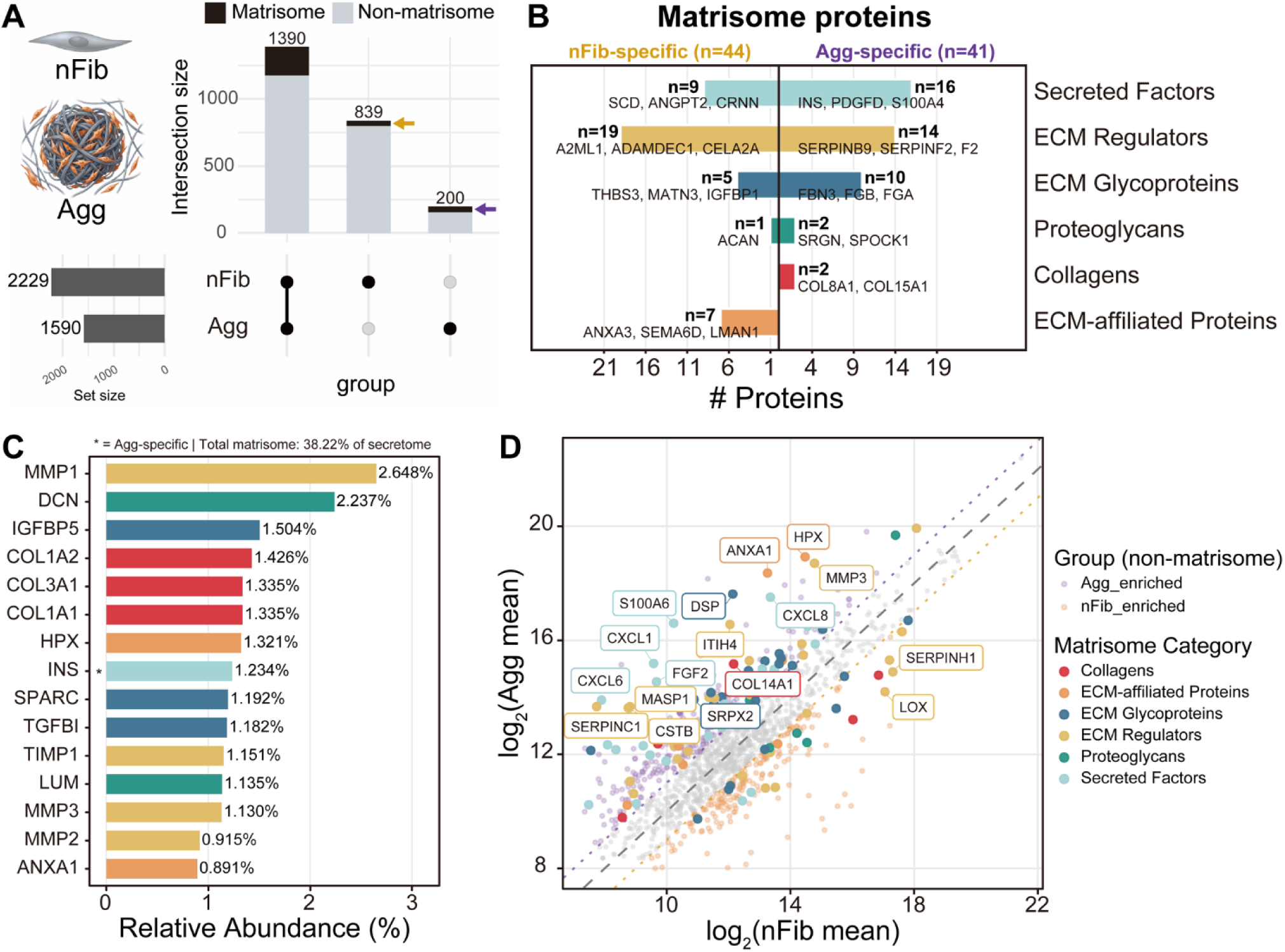
Secretome proteomics reveals a matrisome-enriched extracellular matrix remodeling program in mechanoadaptive fibroblast aggregates. (A) UpSet plot showing overlap of proteins reproducibly identified (≥2 of 3 biological replicates) in aggregate (Agg) and normal fibroblast (nFib) secretomes. Total detected proteins are indicated (Agg, *n*=1,590; nFib, *n*=2,229). Intersection bars denote Agg-specific (*n*=200), shared (*n*=1,390), and nFib-specific (*n*=839) proteins, with matrisome and non-matrisome components shown in black and gray, respectively. Arrows highlight the enrichment of matrisome proteins within Agg- and nFib-specific subsets. (B) Diverging bar plot of matrisome proteins uniquely detected in Agg (*n*=41) or nFib (*n*=44) secretomes. Proteins are grouped by matrisome class, and the three most abundant proteins in each category are annotated. (C) Top 15 matrisome proteins ranked by relative abundance in the Agg secretome, expressed as the percentage of total secretome signal. Matrisome proteins collectively account for 38.22% of the Agg secretome. Asterisks indicate Agg-specific proteins absent from nFib secretomes. (D) Scatter plot comparing mean log_2_ LFQ intensities of proteins detected in both Agg and nFib secretomes. Shared proteins with <2-fold enrichment are shown in grey, whereas non-matrisome and matrisome proteins enriched ≥2-fold in Agg or nFib are highlighted by colored symbols according to condition and matrisome category. Dashed and dotted lines indicate fold-change thresholds of 1 and 2, respectively. Selected matrisome proteins exhibiting high abundance and enrichment in Agg secretomes are labeled. Fold-change values are shown for exploratory comparison.

Ranking the top 15 matrisome proteins by relative abundance in the Agg secretome revealed a coordinated remodeling signature dominated by proteolytic and structural components (Figure 5C). The matrix metalloproteinase 1 (MMP1) was the most abundant matrisome protein (2.648%), co-detected with MMP3 (1.130%) and MMP2 (0.915%), as well as its inhibitor, tissue inhibitor of metalloproteinase 1 (TIMP1; 1.151%), a stoichiometry indicating active, counter-regulated matrix turnover rather than unopposed proteolysis. The decorin (DCN), a proteoglycan (2.237%) and fibrillar collagens COL1A2, COL3A1, and COL1A1 (1.335-1.426%) were among the most abundant structural components. Insulin-like growth factor-binding protein 5 (IGFBP5; 1.504%), secreted protein acidic and rich in cysteine (SPARC) encoding osteonectin (1.192%), and transforming growth factor, beta-induced, 68 kDa (TGFBI; 1.182%) were also among the top abundant ECM glycoproteins.

Fold-change comparison of aggregates and nFib mean label-free quantification (LFQ) intensities identified proteins differentially enriched in either condition (Figure 5D). Among aggregate-enriched matrisome proteins, the Secreted Factors (CXCL8, CXCL1, CXCL6, FGF2, and S100A6) were notably elevated, pointing to an active paracrine signaling environment. ECM Regulators (MMP3, MASP1), ECM Glycoproteins (SRPX2, DSP), Collagen (COL14A1), and ECM-affiliated Proteins (ANXA1, HPX) were also enriched in aggregates. Conversely, the ECM Regulators lysyl oxidase (LOX) and serpin family H member 1 (SERPINH1) were preferentially enriched in the nFib secretome. Collectively, these findings indicate that the Agg secretome is defined by a dual program of active ECM remodeling and paracrine signaling.

## 3. Discussion

We previously demonstrated that the mechanodynamic Gut-on-a-chip platform recapitulates epithelial barrier dysfunction, in which chronic luminal fluid shear stress drives an irreversible transition of shear-vulnerable normal intestinal fibroblasts into shear-resistant, mechanoadaptive fibroblasts [4]. In parallel, mesofluidic and macroscale dynamic culture systems, including rotary shaking, provide complementary platforms for applying sustained mechanical stimulation and evaluating the robustness of fibroblast mechanotransduction across distinct experimental settings. Importantly, these platforms consistently reproduced the identical phenotypic responses in primary intestinal fibroblasts isolated from multiple anatomical regions (small and large intestine) and from independent tissue donors, indicating that the observed mechanoadaptive transition is not donor-or tissue-specific but instead represents a conserved cellular response to chronic fluid shear stress.

The concordance of these independent culture platforms establishes strong reproducibility and generalizability of mechanically induced fibroblast activation across diverse biological sources while minimizing platform-specific bias. Such cross-platform validation is particularly important for studies employing primary human cells, where donor-to-donor variability can confound biological interpretation. Collectively, these complementary systems provide a robust experimental framework for directly interrogating how sustained mechanical forces regulate fibroblast phenotype, mechanoadaptation, and ECM remodeling independent from inflammatory or biochemical stimuli.

Ultrastructural analysis revealed that the fibroblast aggregates comprised a heterogeneous population of both intact and degenerating cells, indicating that they represent a dynamic multicellular microenvironment rather than a random accumulation of cells. Approximately half of the cells retained elongated mitochondria with well-preserved cristae, suggesting that a substantial fraction of the aggregate remained metabolically intact despite its dense microarchitecture. Many cells also exhibited a transition from an elongated spindle-shaped morphology to a stellate, myofibroblast-like phenotype characterized by subplasmalemmal protrusions and slender organelle-poor cellular processes. These features closely resemble the ultrastructure of intestinal subepithelial myofibroblast-like cells and are consistent with cellular activation in response to sustained mechanical stimulation [10]. Extensive interdigitation of these cellular processes generated a continuous intercellular network that is likely to strengthen mechanical coupling while facilitating cell-cell communication throughout the aggregate. The 2D SEM and 3D SBF-SEM further demonstrated that the aggregates were embedded within a dense extracellular fibrillar network. Rather than existing isolated fibers, the fibrils formed a continuous meshwork in which individual strands overlapped, intertwined, and merged into thicker bundles. These fibrillar bundles frequently connected neighboring cells, producing a highly integrated extracellular scaffold with minimal exposed cell surface. The irregular nodular surface, together with the predominance of interconnected fibrils and the absence of abundant free-ending fibers on the exposed cell surface, is consistent with a mature ECM rather than an early fibrillogenic state. Collectively, these findings indicate that chronic mechanical stimulation promotes coordinated remodeling of both the cellular and extracellular compartments, resulting in a mechanically integrated fibroblast aggregate with features characteristic of activated myofibroblasts and a stabilized ECM. This tissue-like architecture provides a physiologically relevant platform for investigating matrix remodeling and mechanobiological processes associated with intestinal fibrosis.

In this ultrastructural context, three prominent features emerged. First, SBF-SEM identified an interconnected network of extracellular void spaces throughout the aggregates. These channels likely facilitate diffusion of nutrients and dissolved oxygen into the aggregate core, while enabling efficient removal of metabolic waste, thereby supporting the viability of interior cells despite the densely packed architecture. Their reproducible presence suggests that they represent an organized structural feature rather than a consequence of random aggregate assembly. Second, these extracellular spaces were enclosed within a highly organized fibrillar microarchitecture extending across both the aggregate surface and the intercellular compartment, a finding consistently observed by both SEM and SBF-SEM. The aggregate surface was almost entirely enveloped by a continuous, interwoven fibrillar meshwork composed of overlapping, braided, and bundled fibers that bridged adjacent cells and matrix domains. In contrast to the short microvilli-like protrusions decorating exposed cell surfaces, the intercellular fibrils formed much longer interconnected networks, likely reinforcing mechanical cohesion and protecting the aggregate against sustained fluid shear. The observed fibrillar organization is consistent with active fibronectin fibrillogenesis, an integrin- and actomyosin-dependent process in which nascent protofibrils elongate and progressively assemble into thicker bundles [11, 12]. Accordingly, the measured fibril diameters (∼110-140 nm) most likely represent bundles rather than individual fibers, consistent with previous descriptions of cell-derived collagen matrices [13]. Rounded surface protrusions measuring approximately 100-300 nm were morphologically consistent with filopodia or clustered microvillus-like structures associated with activated myofibroblasts. This interpretation is supported by the established ultrastructural hallmarks of fibroblast-to-myofibroblast transition, including α-SMA-stress fiber overlapping [14] and fibronexus-mediated coupling between intracellular actin bundles and extracellular fibronectin fibers [15, 16]. Fluid shear stress is a well-recognized inducer of this mechanotransductive program through TGF-β/Rho-ROCK signaling, promoting stress fiber formation and fibroblast alignment along the direction of flow [17–19]. These observations closely mirror our previous Gut-on-a-chip study, in which prolonged fluidic stimulation induced compact 3D fibroblast aggregates with marked increases in α-SMA and collagen I expression together with extensive collagen-like fibrillar deposition [4]. The concomitant elevation of pan-MMP activity further supports the conclusion that these aggregates represent a mechanoadaptive, profibrotic phenotype driven by chronic mechanical stimulation. Third, nearly all cells contained reproducible populations of membrane-bound electron-dense bodies measuring <1 µm in diameter. These structures were predominantly localized in the basal or perinuclear cytoplasm and, in a subset of cells, immediately beneath the plasma membrane. Their size, morphology, and intracellular distribution are consistent with lysosome-related or autophagolysosomal compartments, including lipofuscin-like residual bodies [20], supporting active intracellular turnover during aggregate remodeling [21]. In parallel, slender organelle-poor cellular processes extensively interdigitated with neighboring cells to generate a continuous throat-and-pore intercellular channel network [22]. Together with the surrounding extracellular fibrillar scaffold, this interconnected architecture likely provides the structural basis for mechanical integration of the aggregate while maintaining an open extracellular transport network. These observations support a model in which coordinated cellular interdigitation and ECM assembly collectively establish the mechanically resilient 3D microarchitecture of shear-induced fibroblast aggregates.

Comparative matrisome profiling of mesofluidically stimulated fibroblast aggregates and static fibroblast monolayers revealed a coordinated ECM remodeling program, indicating that mesofluidic stimulation recapitulates key molecular features of early fibrogenic activation. Aggregate secretomes were enriched in fibrillar collagens (COL1A1, COL1A2, and COL3A1), together with the small leucine-rich proteoglycans (SLRPs) such as decorin (DCN) and lumican (LUM), and the fibril-associated collagen COL14A1, which cooperatively regulate collagen fibrillogenesis by restricting fibril diameter and increasing interfibrillar spacing [23]. LUM has also been implicated in promoting fibroblast-to-myofibroblast transition through induction of α-SMA, collagen I, and TGF-β signaling [24]. These molecular changes are consistent with the altered collagen fibril architecture observed by SEM and SBF-SEM, linking matrix composition with ultrastructural remodeling. The aggregates also displayed coordinated enrichment of MMP1, MMP2, MMP3, and their endogenous inhibitor TIMP1, consistent with the dysregulated ECM turnover that characterizes intestinal fibrosis rather than simple matrix accumulation or degradation [25]. Elevated TIMP1 has been associated with increased matrix stiffness in CD strictures [26], whereas reduced net MMP activity distinguishes fibrostenotic CD from ulcerative colitis (UC) [25]. In parallel, enrichment of platelet-derived growth factor D (PDGFD), fibrinogen chains (FGA and FGB), prothrombin (F2), and SERPINF2 suggests activation of a provisional wound-healing matrix coupled with autocrine profibrotic signaling, consistent with established roles of PDGF signaling in fibroblast activation and fibrosis [27]. Beyond structural remodeling, the aggregate secretome exhibited a distinct mechanoadaptive signaling profile characterized by enrichment of chemokines (CXCL8, CXCL1, and CXCL6), together with FGF2 and S100 calcium-binding protein A4 (S100A4). CXCL8 and related chemokines promote fibroblast-to-myofibroblast transition independently of exogenous TGF-β [28], while CXCL1 and CXCL6 have likewise been implicated in fibroblast activation and tissue remodeling [29, 30]. In contrast, FGF2 antagonizes TGF-β-driven myofibroblast differentiation, and S100A6 regulates mechanically induced cytoskeletal remodeling and proliferation [31, 32]. Their concurrent enrichment suggests that the aggregates represent an actively remodeling mechanoadaptive state, in which profibrotic signaling is balanced by adaptive and regenerative pathways. This dynamic signaling architecture may provide a more informative pharmacological endpoint than collagen production alone for evaluating candidate antifibrotic therapies. Additional aggregate-specific proteins further support activation of fibrosis-associated remodeling pathways. MASP1, SRGN, SPOCK1, and COL8A1 collectively implicate complement-coagulation signaling, pericellular ECM remodeling, and TGF-β-associated myofibroblast activation [33–36], whereas ANXA1 and HPX indicate concomitant activation of injury-response and cytoprotective pathways [37, 38]. Together, these findings demonstrate that sustained mechanical stimulation elicits a complex secretory program extending beyond ECM deposition to encompass inflammatory, reparative, and stress-response signaling. In contrast, static fibroblast monolayers were relatively enriched in lysyl oxidase (LOX) and the collagen-specific chaperone SERPINH1 (HSP47), two key regulators of collagen maturation and crosslinking [39, 40]. Their preferential abundance under static conditions (i.e., 2D monolayers) suggests that mesofluidic stimulation redirects fibroblast activity away from constitutive collagen processing toward dynamic ECM remodeling, fibrillogenesis, and paracrine signaling.

Rather than simply increasing matrix production and accumulation, mechanoadaptive aggregates reproduce a regulator-driven remodeling phenotype that closely resembles fibrostenotic intestinal fibrosis. Collectively, these findings demonstrate that mesofluidically generated fibroblast aggregates recapitulate multiple molecular hallmarks of intestinal fibrogenesis, including coordinated collagen remodeling, dysregulated ECM turnover, and mechanically responsive paracrine signaling. Together with the ultrastructural features identified by SEM and volumetric SBF-SEM, these data establish the platform as a physiologically relevant and scalable model for investigating fibrosis mechanisms and for preclinical screening of antifibrotic therapeutics.

Beyond serving as an independent validation of our previous observations, this study substantially expands the biological and translational significance of mechanically induced fibroblast aggregation by defining its ultrastructural and matrisome architecture at high resolution. Integrating SBF-SEM with quantitative proteomics establishes that sustained fluid shear is necessary and sufficient to drive coordinated fibroblast activation, ECM remodeling, and tissue-scale architectural reorganization in the absence of inflammatory or immune cell-derived stimuli. These findings strengthen the concept that aberrant mechanobiological cues are not merely secondary consequences of fibrosis, but may function as primary pathophysiological drivers capable of initiating and sustaining fibrogenic microenvironment. Our previous work demonstrated that these mechanically induced fibroblast aggregates exhibit tissue stiffness comparable to that of human CD strictures [4]. Based on this observation, we propose a mechanobiological model in which chronic epithelial barrier dysfunction exposes subepithelial fibroblasts to abnormal luminal fluid shear, promoting the emergence of shear-resistant, mechanoadaptive fibroblasts. These cells subsequently undergo persistent myofibroblast activation and excessive ECM deposition, progressively generating a mechanically stiffened microenvironment that further reinforces profibrotic signaling through positive mechanotransductive feedback. Such a feed-forward mechanism provides a plausible explanation for the self-perpetuating nature of intestinal fibrosis and offers a novel conceptual framework linking epithelial barrier failure, altered mechanical forces, fibroblast adaptation, and pathological matrix remodeling during fibrostenotic disease progression.

From a translational perspective, the ability to reproducibly generate mechanically activated fibroblast aggregates *in vitro* provides an experimentally tractable model of the fibrotic stromal microenvironment. Unlike conventional static cultures, this platform recapitulates coordinated ultrastructural, biomechanical, and proteomic features associated with fibrotic remodeling under defined mechanical stimulation. To our knowledge, no quantitative *in vitro* model currently captures the structural and functional complexity of intestinal fibrosis. This system therefore offers a scalable platform for dissecting fibrosis mechanisms, identifying mechanotransduction pathways, and evaluating candidate anti-fibrotic therapies under physiologically relevant mechanical conditions.

Because the model is compatible with primary human fibroblasts derived from multiple donors and distinct intestinal regions, it also enables systematic investigation of inter- and intra-patient heterogeneity while maintaining precise control over key experimental variables, including fluid flow, mechanical stimulation, and cellular composition. Together, these features establish a versatile human platform for studying stromal remodeling and support its future application in patient stratification and the development of personalized anti-fibrotic therapies.

One of the major challenges in studying intestinal fibrosis is disentangling the contributions of mechanical forces from the complex inflammatory, immune, microbial, and genetic factors present *in vivo*. Although inflammation is widely recognized as a key driver of CD, increasing evidence indicates that fibrosis can persist or even progress despite effective suppression of inflammatory activity, suggesting that mechanobiological processes become partially autonomous during disease evolution. Human clinical specimens and animal models inherently integrate numerous interacting variables, making it difficult to determine the independent contribution of individual biomechanical cues. In this context, New Approach Methodologies (NAMs) and human microphysiological systems (MPS) provide a unique experimental advantage by enabling precise control of fluid shear stress, tissue deformation, and extracellular mechanical environments while preserving essential features of human intestinal cell biology. Such reductionist yet physiologically relevant platforms allow causal interrogation of mechanotransduction pathways that would otherwise remain obscured within the complexity of native tissues.

Despite these advances, several limitations should be acknowledged. First, the present study establishes proof-of-principle using representative primary intestinal fibroblast populations, but broader validation across larger and more diverse patient cohorts, including fibroblasts isolated from inflammatory, stricturing, penetrating, and healthy intestinal tissues, will be required to establish the generalizability of the mechanoadaptive phenotype. Second, the current model intentionally isolates epithelial and other surrounding cell types and therefore lacks the multicellular complexity of the intestinal mucosa. Incorporating homologous or heterologous patient-derived epithelial cells would enable direct investigation of epithelial-stromal mechanobiological crosstalk under both physiological and diseased conditions. Additional integration of immune cells, smooth muscle cells, or patient-specific microbiota would further recapitulate the inflammatory and microbial microenvironment that shapes fibrosis progression *in vivo*. Finally, combining these multicellular models with longitudinal functional measurements, spatial transcriptomics, and single-cell multi-omics will provide a more comprehensive understanding of how chronic mechanical stimulation remodels the intestinal stromal niche over time.

In summary, this work establishes a mechanically defined human fibrosis model that bridges ultrastructural architecture, proteomics-supported molecular phenotyping, and molecular ECM remodeling in various experimental platforms. By demonstrating that sustained fluid shear alone is sufficient to induce persistent fibroblast mechanoadaptation and fibrosis-associated matrix remodeling, our findings support the emerging paradigm that aberrant biomechanics constitute an independent pathogenic axis in intestinal fibrosis. Beyond providing new insight into the pathobiology of fibrostenotic CD, this platform offers a robust and scalable foundation for mechanistic studies, biomarker discovery, and preclinical evaluation of anti-fibrotic therapies in a human-relevant experimental system.

## 4. Experimental Section

### 4.1. Device Microfabrication

The microfluidic Gut-on-a-chip device was fabricated using standard soft lithography as previously described [5, 8]. Briefly, a degassed 10:1 (wt wt^-1^) mixture of polydimethylsiloxane (PDMS; Sylgard 184, Dow Corning) base polymer and curing agent was cast onto the silanized silicon master containing SU-8-defined microfluidic features and cured at 60 °C for 6 h. The cured PDMS was peeled from the mold, trimmed, and punched to create inlet and outlet ports before device assembly. The upper (500 μm height) and the lower (200 μm height) microchannel layers were irreversibly bonded to a porous PDMS membrane (10 μm pore diameter, 25 μm center-to-center spacing, 10 μm thickness) [41] following oxygen plasma activation (COVANCE-1MPR, Femto Science Inc.), then incubated at 80 °C for 6 h. Each microchannel was connected to silicone tubing (Tygon 3350, Beaverton) via bent blunt-end 18-gauge stainless needles (Kimble Chase) to enable cell seeding and continuous perfusion of culture medium into microchannels.

### 4.2. Cell Culture

Primary human intestinal fibroblasts from the small and large intestine were used to investigate shear-induced aggregate formation. Normal small intestinal fibroblasts (cat. no. 2920; ScienCell), normal colonic fibroblasts (cat. no. 2880; ScienCell), and patient-derived colonic fibroblasts isolated from histologically normal colonic tissue were cultured in Dulbecco’s Modified Eagle Medium (DMEM, cat. no. 11995073; Gibco) supplemented with 20% (vol vol^-1^) heat-inactivated fetal bovine serum (FBS, cat. no. A5256502; Gibco), 1% (wt vol^-1^) L-glutamine (cat. no. 25-030-149; Gibco), and antibiotics (100 U mL^-1^ Penicillin and 100 µg mL^-1^ Streptomycin, cat. no. 10-378-016; Gibco). Cells (passage number <6) were maintained in T75 flasks at 37 °C in a humidified atmosphere containing 5% CO_2_ and passed at approximately 80% confluence.

For the isolation of donor-derived colonic fibroblasts, de-identified colonic biopsy (4-5 fragments, ∼2 mm^2^ each) were obtained from a normal donor under an approved protocol by the Institutional Review Board (IRB; 06-050) of Cleveland Clinic Tissue Center. A consent was not required for this specimen, as the tissue was procured from discarded surgical material in accordance with IRB guidelines. The specimen was washed twice with Ca^2+^- and Mg^2+^-free Hank’s Balanced Salt Solution (HBSS, cat. no. MT21023CV; Corning) and enzymatically digested in HBSS containing collagenase types I, II, and IV (100 U mL^-1^ each, cat. no. C1639, C1764, and C5138, respectively; MilliporeSigma) at 37 °C for 45-60 min with continuous agitation (140 rpm). DNase (100 U mL^-1^ final concentration; cat. no., LS002139, Worthington) was subsequently added to dissociate DNA-associated cell aggregates, followed by an additional 30 min incubation at 37 °C with gentle agitation (60 rpm). The suspension was filtered through a 70 μm cell strainer (cat. no. 229483; Celltreat Scientific Products), and centrifuged (300× *g*, 10 min). The resulting cell pellet was washed twice with Ca^2+^- and Mg^2+^-free HBSS, resuspended in a growth medium, and seeded into 6-well plates. Cultures were maintained at 37 °C with medium changes every other day until reaching approximately 80% confluence.

### 4.3. Induction of Fibroblast Aggregates

Fibroblast aggregates were generated using either a microfluidic Gut-on-a-Chip platform or a mesofluidic shaker-based culture system. For microfluidic culture, fibroblasts were seeded into the upper microchannel of Gut-on-a-Chip devices coated with Matrigel (0.3 mg mL^-1^, cat. no. CB-40234; Corning) and collagen I (0.03 mg mL^-1^, cat. no. A10483-01; Gibco) following ultraviolet/ozone surface activation (UVO Cleaner 32, Jelight Company Inc.). Cells were allowed to attach under static conditions for 1.5 h before continuous perfusion with culture medium at 30 µL h^-1^ for up to 160 h. Under these conditions, sustained luminal shear stress (∼0.0013 dyne cm^-2^) reproducibly induced spontaneous formation of three-dimensional fibroblast aggregates, as previously reported [4]. For mesofluidic culture, fibroblasts were seeded onto identically coated porous PDMS membranes mounted in 6-well plates and cultured overnight under static conditions to establish confluent monolayers. The plates were subsequently transferred to an orbital shaker and continuously rotated at 75 rpm to generate sustained fluid shear stress. Progressive detachment of the fibroblast monolayer was followed by spontaneous self-assembly into multicellular aggregates. Aggregate formation was monitored by phase-contrast microscopy, and experiments were terminated after reproducible aggregate formation, typically at approximately 48 h.

### 4.4. Measurement of Matrix Metalloproteinase Activity

Extracellular pan-matrix metalloproteinase (pan-MMP) activity was quantified using a fluorometric MMP Activity Assay Kit (Abcam; cat. no. ab112146), which detects the combined activity of multiple MMPs, including MMP-1, -2, -3, -7, -8, -9, -10, -12, -13, and -14. Culture supernatants collected from aggregate and non-aggregate conditions (i.e., monolayer cultures) were centrifuged at 1,000× *g* for 10 min to remove cellular debris. Clarified supernatants were mixed 1:1 (vol vol^-1^) with 2 mM 4-aminophenylmercuric acetate to activate latent MMPs and incubated at 37 °C for 15 min.

Fluorogenic pan-MMP substrate was then added according to the manufacturer’s instructions, and fluorescence was recorded every 10 min for 90 min at 37 °C using a microplate reader (BioTek Instruments) with excitation and emission wavelengths of 490 and 525 nm, respectively. All samples were analyzed in technical duplicates, and data were averaged from at least three independent biological replicates.

### 4.5. Morphological Analysis

Fibroblast monolayers and aggregates were monitored by phase-contrast microscopy using an inverted microscope (DMi1, Leica Microsystems) equipped with 10× (NA 0.22) and 20× (NA 0.30) objectives, a digital camera (MC120 HD, Leica Microsystems), and Leica Application Suite software (LAS v4.12, Leica). For each condition and time point, images were acquired from at least 10 randomly selected fields of view across a minimum of two independent biological replicates. Representative images are shown.

For immunofluorescence analysis, samples were fixed with 4% (wt vol^-1^) paraformaldehyde (PFA; cat. no. 15710; Electron Microscopy Sciences) for 30 min, permeabilized with 0.3% (vol vol^-1^) Triton X-100 (MilliporeSigma) for 15 min, and blocked with 2% (wt vol^-1^) bovine serum albumin (BSA; MilliporeSigma) in phosphate-buffered saline (PBS; Gibco) for 1 h at room temperature. PBS washes were performed between each step. Samples were incubated with primary antibodies against α-SMA (mouse, cat. no. ab7817; Abcam) and COL I (rabbit, cat. no. ab90395; Abcam) for 1 h at room temperature, followed by Alexa Fluor 488-conjugated donkey anti-mouse (cat. no. ab150105; Abcam) and Alexa Fluor 555-conjugated goat anti-rabbit (cat. no. ab150078; Abcam) secondary antibodies for 1 h in the dark. Nuclei and filamentous actin (F-actin) were counterstained with 4’,6-diamidino-2-phenylindole dihydrochloride (DAPI, 1 μg mL^-1^, cat. no. 62248; Thermo Fisher) and CruzFluor 647-conjugated phalloidin (1:500, cat. no. sc-363797; Santa Cruz Biotechnology), respectively. Samples were mounted with Fluoromount mounting medium (MilliporeSigma).

Confocal images were acquired using a Leica TCS SP8 laser-scanning confocal microscope equipped with a 25× water-immersion objective (NA 0.95), 405-, 488-, 561-, and 633-nm excitation lasers, and hybrid (HyD) detectors. Single optical sections and Z-stack images were acquired using Leica LAS X software. Orthogonal reconstructions of Z-stack images were generated to visualize vertical cross-sectional architecture. Fluorescence intensity was quantified from three randomly selected regions of interest per condition, and cell layer thickness was measured from 10 independent cross-sectional images using ImageJ (NIH).

### 4.6. Ultrastructural Analysis

For scanning electron microscopy (SEM), fibroblast aggregates and static monolayers were fixed in 4% (wt vol^-1^) PFA and 2.5% (wt vol^-1^) glutaraldehyde (cat. no. 16320; Electron Microscopy Sciences) in PBS for 1 h at room temperature, followed by post-fixation with 1% (wt vol^-1^) osmium tetroxide in 0.1 M sodium cacodylate buffer (cat. no. 11650; Electron Microscopy Sciences) for 1 h at 4 °C. Samples were rinsed with PBS and dehydrated through a graded ethanol series (30%, 50%, 70%, 80%, 90%, 95%, and 100%; 10 min per step), followed by treatment with hexamethyldisilazane (HMDS; cat. no. 16700; Electron Microscopy Sciences) for 10 min. Samples were air-dried overnight in a vacuum desiccator containing anhydrous desiccant (cat. no. 23005; Drierite) before being mounted on aluminum stubs and sputter-coated with ∼10 nm layer of gold. SEM imaging was performed using a Zeiss SIGMA VP Scanning Electron Microscope (Carl Zeiss Inc.).

For serial block-face scanning electron microscopy (SBF-SEM), fibroblast aggregates were fixed in 4% (wt vol^-1^) PFA and 2.5% (wt vol^-1^) glutaraldehyde prepared in 0.1 M sodium cacodylate buffer. The aggregates were processed using an automated tissue staining protocol on an ASP-1000 system (Microscopy Innovations) as previously described [42]. Briefly, samples were sequentially stained with reduced osmium (2% wt vol^-1^ osmium tetroxide and 2% wt vol^-1^ potassium ferrocyanide), 1% (wt vol^-1^) thiocarbohydrazide, 2% (wt vol^-1^) osmium tetroxide, 1% (wt vol^-1^) aqueous uranyl acetate, and Walton’s lead aspartate. Following heavy-metal staining, samples were dehydrated through a graded ethanol series, transitioned through propylene oxide, and embedded in medium-hardness Embed 812 epoxy resin (Electron Microscopy Sciences). Polymerized resin blocks were trimmed, mounted on aluminum specimen pins, and coated with colloidal silver to improve electrical conductivity. Serial block-face imaging was performed using either a VolumeScope II system (Thermo Fisher Scientific) or a 3View in-chamber ultramicrotome (Gatan) integrated with a Sigma VP field-emission scanning electron microscope (Zeiss). Images were acquired under high vacuum using a low-kV backscattered electron detector (Gatan) at an accelerating voltage of 2.0 kV with a 30 μm aperture. Image volumes covering approximately 65 × 65 μm were collected at a lateral resolution of 6 nm per pixel, a section thickness of 60 nm, and a total imaging depth of 200-500 serial sections. Image registration, 3D reconstruction, and video generation were performed using FIJI/ImageJ (NIH). The extracellular space was segmented from individual image slices by intensity-based thresholding followed by manual refinement using the FIJI 3D Viewer plugin. Quantitative pore network analysis was subsequently performed in PoreSpy (Python). Automated pore network extraction was carried out using the SNOW2 (Sub-Network of an Over-segmented Watershed 2) algorithm to better represent the anisotropic geometry of intercellular voids within fibroblast aggregates [43], which was modified to calculate ellipsoidal pore dimensions in addition to conventional largest-inscribed-sphere measurements of pores and pore throats.

### 4.7. Sample Preparation and LC-MS/MS Proteomic Analysis

Culture medium was concentrated to approximately 70-75 μL using 3 kDa molecular weight cutoff centrifugal filters. Proteins (45 μg per sample) were processed using S-Trap spin columns (Protifi) according to the manufacturer’s protocol. Briefly, samples were lysed in sodium dodecyl sulfate (SDS)-containing buffer, and protein concentrations were determined by bicinchoninic acid (BCA) assay. Proteins were reduced with dithiothreitol (DTT), alkylated with iodoacetamide (IAA), acidified with phosphoric acid, and diluted with methanol-based binding buffer before loading onto S-Trap columns. Following 4 washes with binding buffer, proteins were digested overnight at 37 °C with sequencing-grade trypsin (1:10, enzyme-to-protein ratio). Peptides were sequentially eluted with 50 mM triethylammonium bicarbonate (TEAB), 0.2% formic acid, and 50% acetonitrile, dried under vacuum, reconstituted in 0.1% formic acid, and filtered through a 0.22 μm membrane before LC– MS/MS analysis. Peptide analysis was performed using a timsTOF Pro 2 quadrupole time-of-flight mass spectrometer (Bruker Daltonics) equipped with a CaptiveSpray ion source and coupled to a nanoLC system. Peptides were separated on a 15 cm × 75 μm internal diameter C18 reversed-phase column (ReproSil AQ, 1.9 μm, 120 Å; Bruker) using a 0.1% formic acid/acetonitrile gradient at a flow rate of 0.3 μL min^-1^. Approximately 1 μL of each sample was injected for analysis. Data were acquired in positive ion mode using a data-dependent acquisition (DDA) workflow with Parallel Accumulation-Serial Fragmentation (PASEF). TIMS-MS survey scans were collected over an m/z range of 100-1,700 and an ion mobility range of 0.60-1.60 Vs cm^-2^, followed by 10 PASEF MS/MS scans per acquisition cycle (1.2 s cycle time). Precursor ions with charge states of 2-5 and intensities above 2,500 arbitrary units were selected for fragmentation, and dynamic exclusion was applied for 0.4 s.

### 4.8. Quantitative Proteomic Analysis

To minimize systematic differences in protein loading between samples, label-free quantification (LFQ) intensities were normalized using a common-protein median normalization approach.

Proteins detected in all six samples (Aggregate (Agg), *n*=3; normal fibroblast (nFib), *n*=3) were used as normalization anchors. For each sample, a scaling factor was calculated as the ratio of the global median intensity (median of the per-sample medians of anchor proteins) to the corresponding sample median. All LFQ intensities were multiplied by the resulting scaling factor before downstream analyses. Zero LFQ values were treated as missing and excluded from normalization and fold-change calculations. Proteins were filtered for reproducibility by retaining those detected (LFQ>0) in at least two of three biological replicates within each condition (Figure S4A). Retained proteins were classified into four mutually exclusive groups according to their detection pattern and relative abundance (Figure S4B): Agg-specific, detected in ≥2/3 Agg replicates and <2/3 nFib replicates; Agg-enriched, detected in both conditions with a mean fold change ≥2 in Agg; nFib-enriched, detected in both conditions with a mean fold change ≥2 in nFib; and Shared, exhibiting <2-fold differences between conditions. Fold changes were calculated as the ratio of mean normalized LFQ intensities (Agg/nFib). Because Agg samples were processed in two independent experimental batches, only one of which included matched nFib samples, fold-change analyses were considered exploratory and were not subjected to formal statistical testing. Matrisome annotation and selection of Agg-enriched matrisome proteins generated the final protein set for downstream analyses (Figure S4B).

Proteins were annotated against the human matrisome using MatrisomeAnalyzeR [44] and MatrisomeDB 2.0 [9]. Annotated proteins were classified as collagens, ECM glycoproteins, proteoglycans (core matrisome), or ECM regulators, ECM-affiliated proteins, and secreted factors (matrisome-associated). Relative protein abundance within the Agg secretome was calculated as the percentage of the mean LFQ intensity of each protein relative to the summed mean LFQ intensity of all detected Agg proteins. Secretome overlap between Agg and nFib samples was visualized using UpSet plots generated with ComplexUpset (v1.3.3), with intersections annotated according to matrisome and non-matrisome composition. Scatter plots comparing mean LFQ intensities between Agg and nFib samples highlighted matrisome proteins exhibiting ≥2-fold differential abundance. Protein labels were assigned using two simultaneous criteria: i) abundance within the top 50th percentile of all detected matrisome proteins in the Agg secretome and ii) fold change within the upper quartile of Agg-enriched proteins or lower quartile of nFib-enriched proteins (log_2_ fold change ≥75th percentile for Agg-enriched proteins or ≤25th percentile for nFib-enriched proteins).

### 4.9. Statistical Analysis

Proteomic data analyses were performed in R (v4.4.2). Data visualization was conducted using the ggplot2 (v4.0.2), patchwork (v1.3.0), and ComplexUpset (v1.3.3) packages, and matrisome annotation was performed using MatrisomeAnalyzeR (v1.0.1). For comparisons of cell layer thickness, α-SMA, COL I, and pan-MMP activity between monolayer and aggregate conditions, statistical significance was assessed using unpaired two-tailed Student’s *t*-tests or Welch’s *t-*test, as appropriate. Data are presented as mean ± SEM unless otherwise indicated, with individual biological replicates shown. A two-sided *p*<0.05 was considered statistically significant. Statistical analyses were performed using GraphPad Prism (v10; GraphPad Software).

## Acknowledgements

This work was supported in part by the NIH NCI IMAT program (1R33CA286797), the Kenneth Rainin Foundation Innovator Award (to H.J.K), the Crohn’s and Colitis Foundation of America Senior Research Award (to H.J.K.), the Bio-industrial Technology Development Program from the Ministry of Trade, Industry & Energy Korea (20018770; to H.J.K. and D.-W.L.), the National Research Foundation of Korea grant funded by the Korean government (MSIT; RS-2025-02215093 to D.-W.L.), and Clinical and Translational Science Collaborative of Northern Ohio funded by the NIH NCATS (UM1TR004528; to H.J.K.). During the preparation of this work, the authors utilized generative artificial intelligence (AI) tools (e.g., Gemini) and illustration platforms (e.g., BioRender.com) in part for the creation of schematics. We appreciate Haerim Jun for assistance with processing movie data.

## Data Availability Statement

The data that support the findings of this study are available from the corresponding author upon reasonable request.

## Author Contributions

S.M. and H.J.K. conceived and designed the study, performed the experiments, analyzed and interpreted the data, and wrote the manuscript. H.S.J. and D.-W.L. performed the proteomic analyses and contributed to data interpretation. G.K. and E.B. conducted the SEM and SBF-SEM experiments and analyzed the ultrastructural data. All authors reviewed and approved the final version of the manuscript.

