## Supplementary text and figures for "Ultrastructural and Proteomic Signatures of Mechanoadaptive Fibroblast Remodeling across Microphysiological and Mesoscale Shear Platforms"

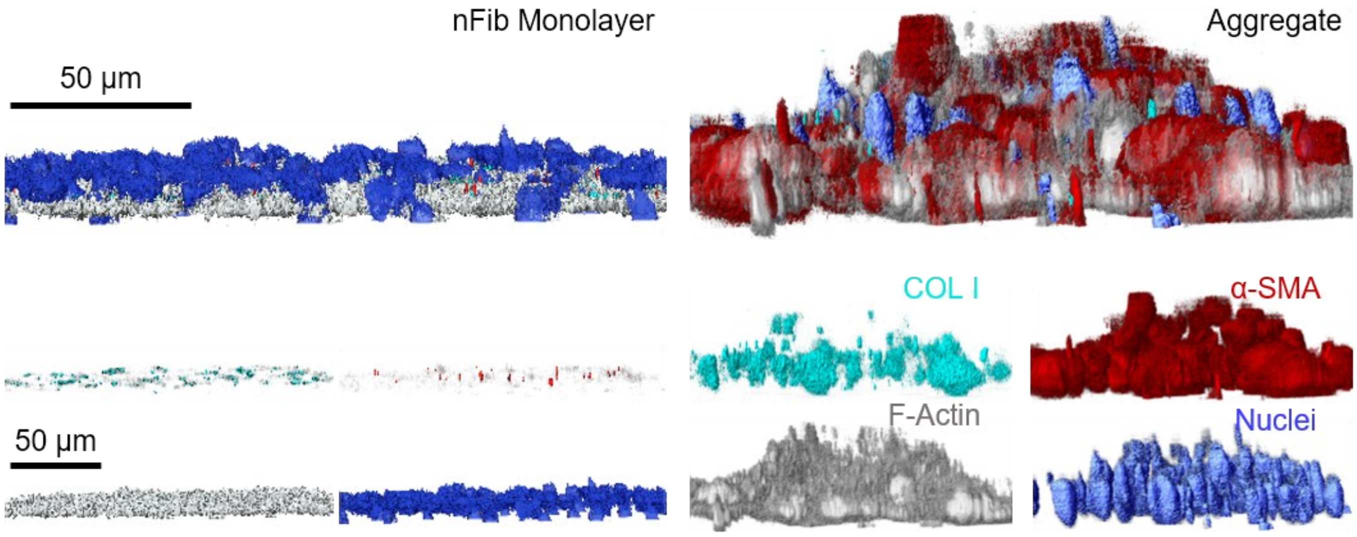


**Figure S1. Individual immunofluorescence channels underlying the composite volumetric reconstruction in Figure 1C.** Merged and single-channel 3D renderings of static nFib monolayer (left) vs. fluidically stimulated aggregate (right), corresponding to the composite image shown in Figure 1C. Top row: merged reconstruction (all channels overlaid). Bottom rows: individual channels shown separately with COL I (cyan), α-SMA (red), F-actin (grey), and nuclei (blue), illustrating the individual contribution of each marker to the composite signal. This figure confirms that the increased height and signal intensity observed in the aggregate (Figure 1C) reflect coordinated upregulation across all channels rather than a single dominant marker.

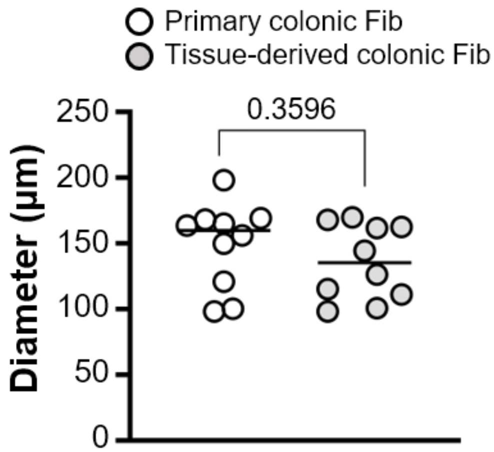


**Figure S2. Aggregate diameter is comparable between primary and tissue-derived colonic fibroblasts under mesofluidic shaking culture.** Scatter plot of aggregate diameter (μm) measured at 48 h for aggregates generated from primary colonic fibroblasts (open circles) versus tissue-derived colonic fibroblasts (grey circles) using the mesofluidic rotary shaker platform. Horizontal lines indicate group means. No significant difference in aggregate diameter was observed between fibroblast sources ($p$=0.3596, Welch's t test), supporting consistent aggregate morphogenesis across independent colonic fibroblast populations. $n$=10 aggregates per group. Statistical significance was determined using Welch's $t$-test.

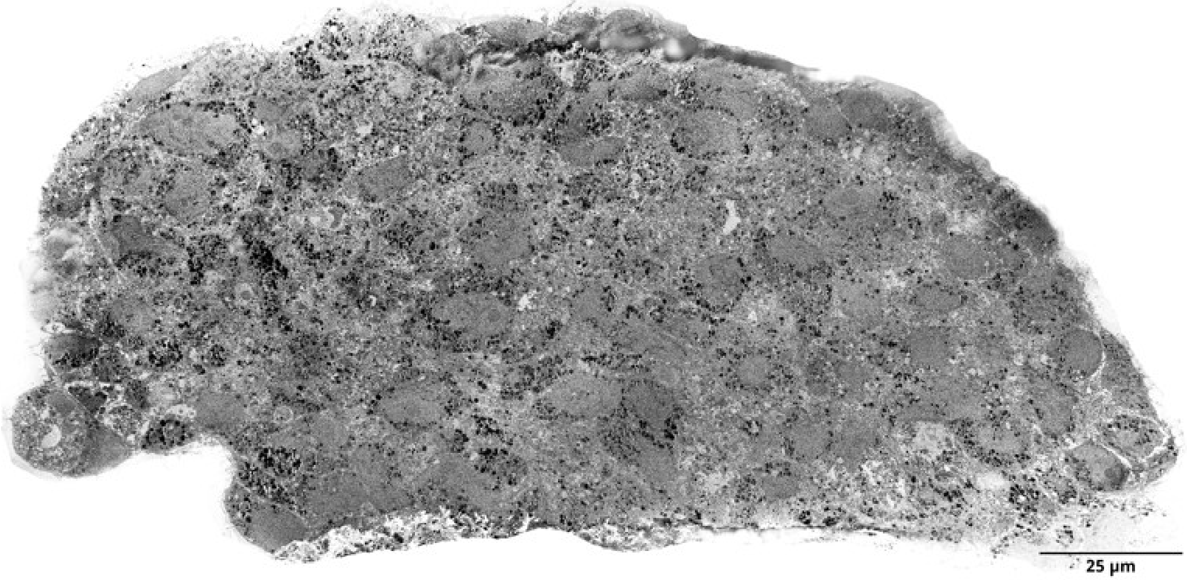


**Figure S3. Whole-aggregate transverse block-face section acquired by serial block-face SEM.** Representative montaged serial block-face scanning electron microscopy (SBF-SEM) image showing the entire transverse cross-section of a mechanoadaptive fibroblast aggregate at a single block-face depth. The montage, assembled from adjacent high-resolution image tiles, reveals the overall multicellular architecture, including densely packed fibroblasts with heterogeneous nuclear morphology, abundant membrane-bound electron-dense cytoplasmic inclusions, and an irregular, lobulated aggregate boundary. This low-magnification overview provides the structural context for the higher-resolution ultrastructural analyses presented in Figure 4.

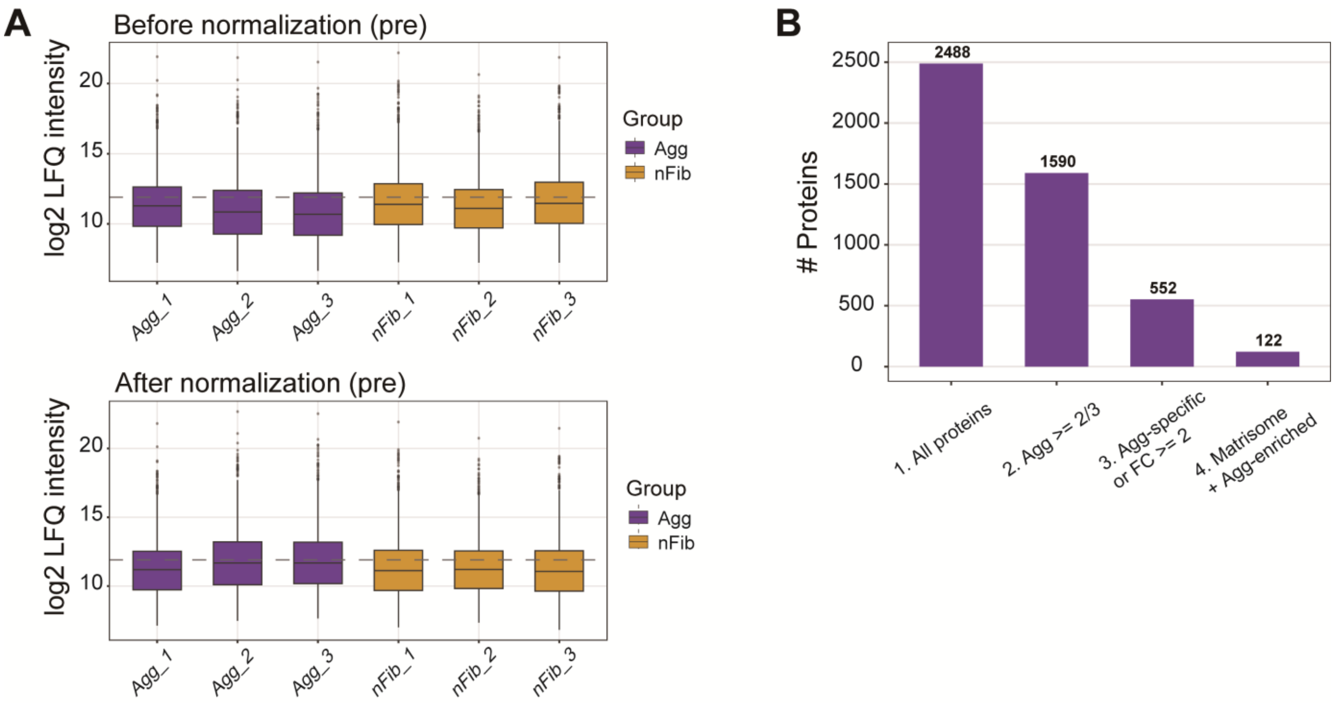


**Figure S4. Proteomic data quality control and sequential filtering pipeline defining the aggregate-associated matrisome subset.**

(A) Boxplots of $log_2$-transformed label-free quantification (LFQ) intensity distributions for each biological replicate (Agg_1-3, purple; nFib_1-3, gold) before (top) and after (bottom) common-protein median normalization. The dashed line indicates the global median intensity across all samples. Normalization equalizes run-level differences in total protein loading across samples while preserving relative intensity distribution of each sample, confirming comparable quantitative dynamic range between the Agg and nFib groups prior to differential analysis. (B) Sequential filtering workflow applied to define the aggregate-associated proteome. Starting from all proteins identified across the dataset (2,488), proteins were retained if reproducibly detected in at least 2 of 3 Agg replicates (1,590), then further restricted to proteins that were either Agg-specific (undetected in nFib) or Agg-enriched (fold change ≥2 relative to nFib) (552). This Agg-enriched/specific subset was finally cross-referenced against matrisome gene ontology annotations to yield the final matrisome-classified, Agg-enriched protein set used for downstream analysis (122).

**Legend for Movies**

**Movie S1.** Rotational 3D volume reconstruction of the aligned SBF-SEM image stack, corresponding to the reconstruction shown in Figure 4B (left panel). The volume was generated by rigid alignment and stacking of sequential block-face section images acquired along the z-axis, preserving native grayscale ultrastructural contrast throughout the reconstructed block. The movie rotates the volume to display the continuous multicellular architecture, including cell boundaries, nuclear and cytoplasmic organelle content, mark matters, and the interdigitating intercellular channel network across all three orthogonal faces of the reconstructed stack. Duration, 30 s.

**Movie S2.** Rotational 3D view of the segmented extracellular space within the SBF-SEM-reconstructed fibroblast aggregate, corresponding to the inset in Figure 4B (middle panel). The extracellular space was isolated by segmentation of the aligned SBF-SEM image stack and rendered as an intensity-coded surface (cyan-to-white scale reflecting local channel width or signal intensity), revealing a continuous, branched network of narrow throat-like segments connecting discrete wider pore-like expansions. The movie rotates the segmented volume to visually display the full 3D connectivity and geometry of the channel network independent of the overlying cellular ultrastructure shown in Movie S1. Duration, 15 s.
